# Similarity of species composition of sand flies (Diptera: Psychodidae) in caves of Quadrilátero Ferrífero, state Minas Gerais, Brasil

**DOI:** 10.1101/703231

**Authors:** Aldenise Martins Campos, Rodrigo dos Anjos Maia, Débora Capucci, Adriano Pereira Paglia, José Dilermando Andrade Filho

## Abstract

Caves, though seemingly inhospitable due to the lack of light in areas furthest from the cave entrance and the low availability and variety of resources, can harbor many species of vertebrates and invertebrates, including phlebotomine sand flies, the vectors of species of parasites of the genus *Leishmania* that cause leishmaniasis. This study aimed to evaluate the species composition of sand flies in located caves at Moeda Sul (MS) and Parque Estadual Serra do Rola Moça (PESRM), in southeastern Brazil at region of the Quadrilátero Ferrífero, state of Minas Gerais. The collections of sand flies were performed with automatic light traps. Were used the Non-Metric Multidimensional Scaling based on dissimilarity matrix calculated with the Jaccard índex and Multivariate Analysis of Permutation to evaluate similarity of the species composition of sand flies among entrance, interior, and surroundings environments of each sampled cave at MS and PESRM, and thus to infer biological mechanisms from the patterns of distribution of sand flies among these different cave environments. A total of 375 phlebotomine sand flies were collected from six genera and 14 species. The most abundant species were *Evandromyia tupynambai* (54.7%), *Brumptomyia troglodytes* (25.6%), *Evandromyia edwardsi,* (6.1%) *Psathyromyia brasiliensis* (4.8%) and *Lutzomyia longipalpis* (4.3%). In place MS, 30 individuals were collected, 16 inside the MS cave and 14 its surroundings. In place PESRM, five individuals were collected in surroundings of RM38 cave; a total of 190 individuals at RM39 cave (48 in the cave and 142 its surroundings) and 150 individuals at RM40 cave, 42 in the cave and 108 its surroundings. Our results showed rich fauna of sand flies with a species composition of sand flies similar among the entrance, interior, and surroundings environments of each sampled cave, suggesting that both caves and their surroundings are important for the maintenance of sand fly communities.

## Introduction

Phlebotomine sand flies are insects of the family Psychodidae, subfamily Phlebotominae (Diptera). They are the invertebrate hosts for protozoan parasites of the genus *Leishmania,* which cause leishmaniases in humans and other mammals [1]. These insects can be found in urban, rural and wild environments, including caves [2, 3, 4, 5, 6, 7, 8, 9, 10].

Cave environments, though seemingly inhospitable due to the lack of light in areas furthest from the cave entrance and the low availability and variety of resources [11, 12], can harbor many species of vertebrates and invertebrates, including phlebotomine sand flies [13, 6, 14, 15, 16, 17, 18, 10].

Studies conducted inside caves have produced interesting results, such as finding that the diversity of sand flies inside caves is greater than or equal to that in their surrounding environments [2, 3, 4, 5, 19, 16]. Additionally, the species composition of sand fly communities of caves differ from that of their surrounding areas with some species being restricted to the caves, other species restricted to surroundings and yet others common to both environments [4, 5, 19, 6, 16, 10]. [10] have also shown that sand flies exhibit differences in their activity rhythms among the cave interior, entrance and surrounding areas due to differences in the presence of sunlight, which suppresses flight activity of these mainly nocturnal insects. However, despite all these investigations, little is known of the patterns of distribution of sand flies in caves and their surroundings, even though it is known that there are different species outside and inside of caves.

Dissimilarity measures have been widely used for evaluating the distribution of different biotas and they have a long-standing history [20, 21, 22]. Thus, this study aimed was to evaluate patterns of similarity in species composition of sand flies among the entrance, interior, and surroundings environments of caves at two different locations at region of the Quadrilátero Ferrífero, state of Minas Gerais. The use of dissimilarity indexes is important for inferring biological mechanisms from the patterns of distribution of sand flies. Although the studied caves are not tourist attractions, the knowledge of the behavior of these insects in the caves and their surroundings is important to for understanding the risk of transmission of *Leishmania* between the sand flies and the vertebrates who visit or live in these environments or near them, including the human population.

## Materials and Methods

### Study area

The study included caves and their surroundings at two different locations in the state of Minas Gerais: Moeda Sul (MS) and Parque Estadual Serra do Rola Moça (PESRM) (Fig 1). These locations are within the region of the Quadrilátero Ferrífero of Minas Gerais, a region known to have a great number of recorded caves, of intense mining activity, exploitation for real estate and urbanization [23, 24]. These region has abundant ferruginous fields called “canga”, a rare geological formation only found in the Quadrilátero Ferrífero of Minas Gerais and in Carajás, in the state of Pará [25, 26], and high altitude fields beyond areas of “capoeira” (secondary formations) [26, 27].

**Fig 1.**
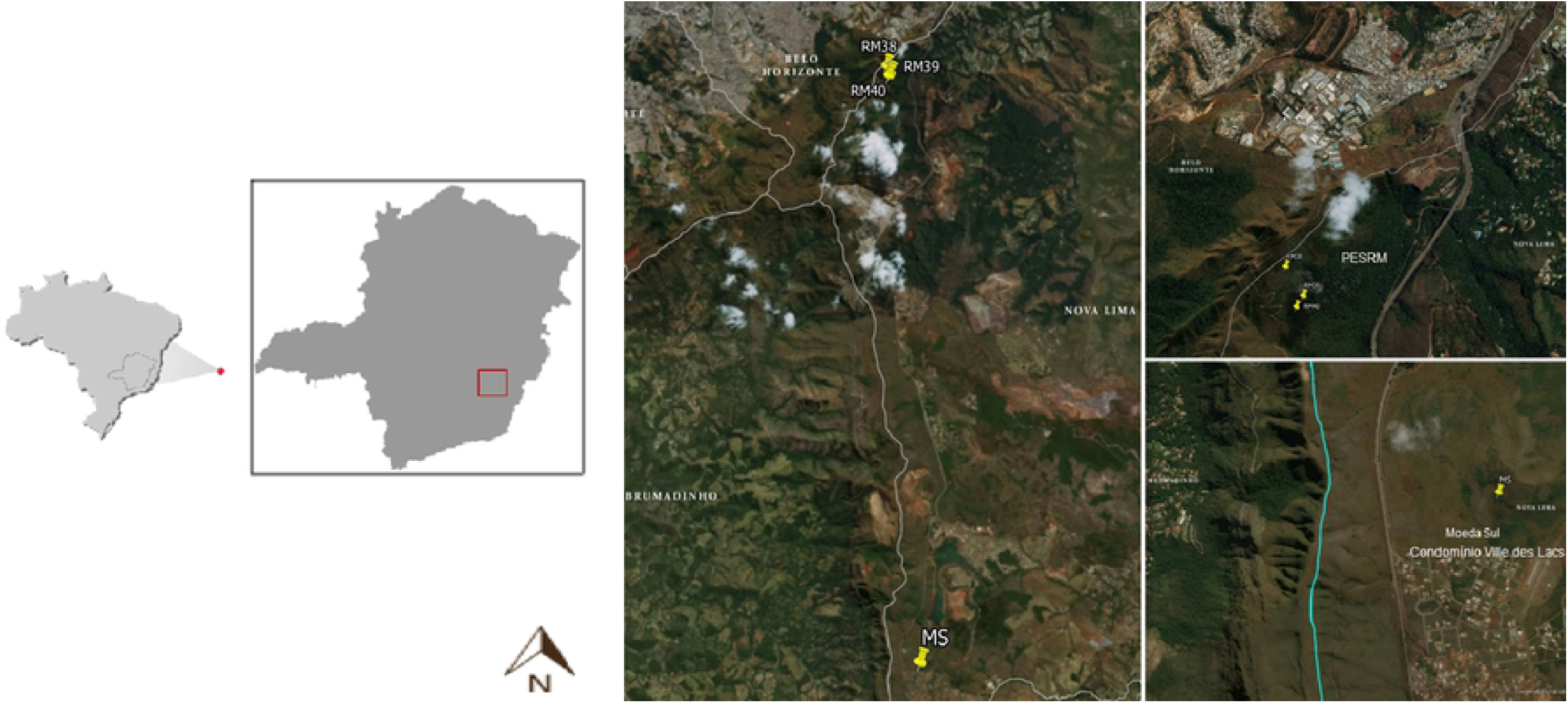
Location of the study area in the Moeda Sul (MS) and Parque Estadual Serra do Rola Moça (PESRM), municipalities of Nova Lima-MG.

The study took place in four caves and their surroundings, one located at MS (MS cave) and three at PESRM (caves RM38, RM39 and RM40), (Fig 2, Fig 3, Fig 4, Fig 5 and Table 1). All the caves were named by Centro Nacional de Pesquisa e Conservação de Cavernas (CECAV). The caves located at PESRM sampled in this study are about 500 m from each other, and about 20 km from the cave located at MS. The caves named RM39 and RM40 do not have an aphotic zone, whereas caves MS and RM38 have aphotic zone about 5 m into the interior from the entrance.

**Fig 2.**
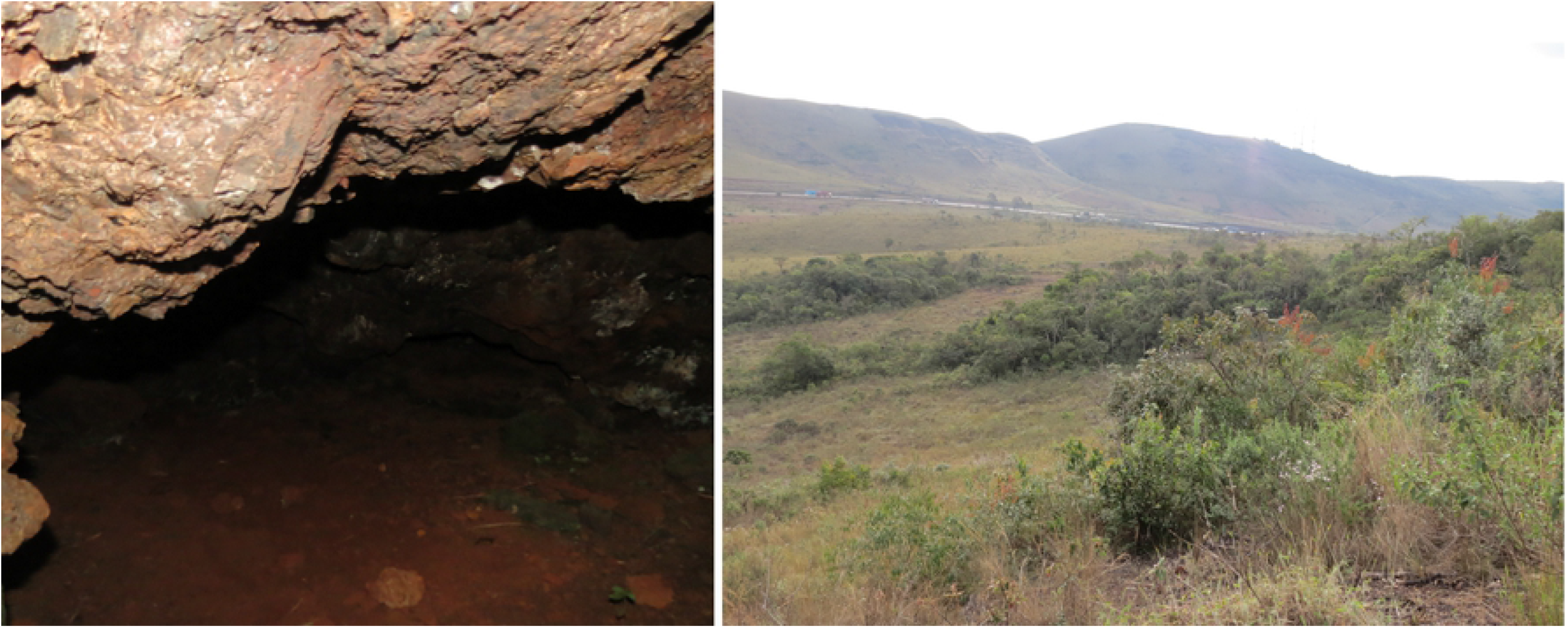
The MS cave and their surroundings in the Moeda Sul (MS), Minas Gerais, Brazil.

**Fig 3.**
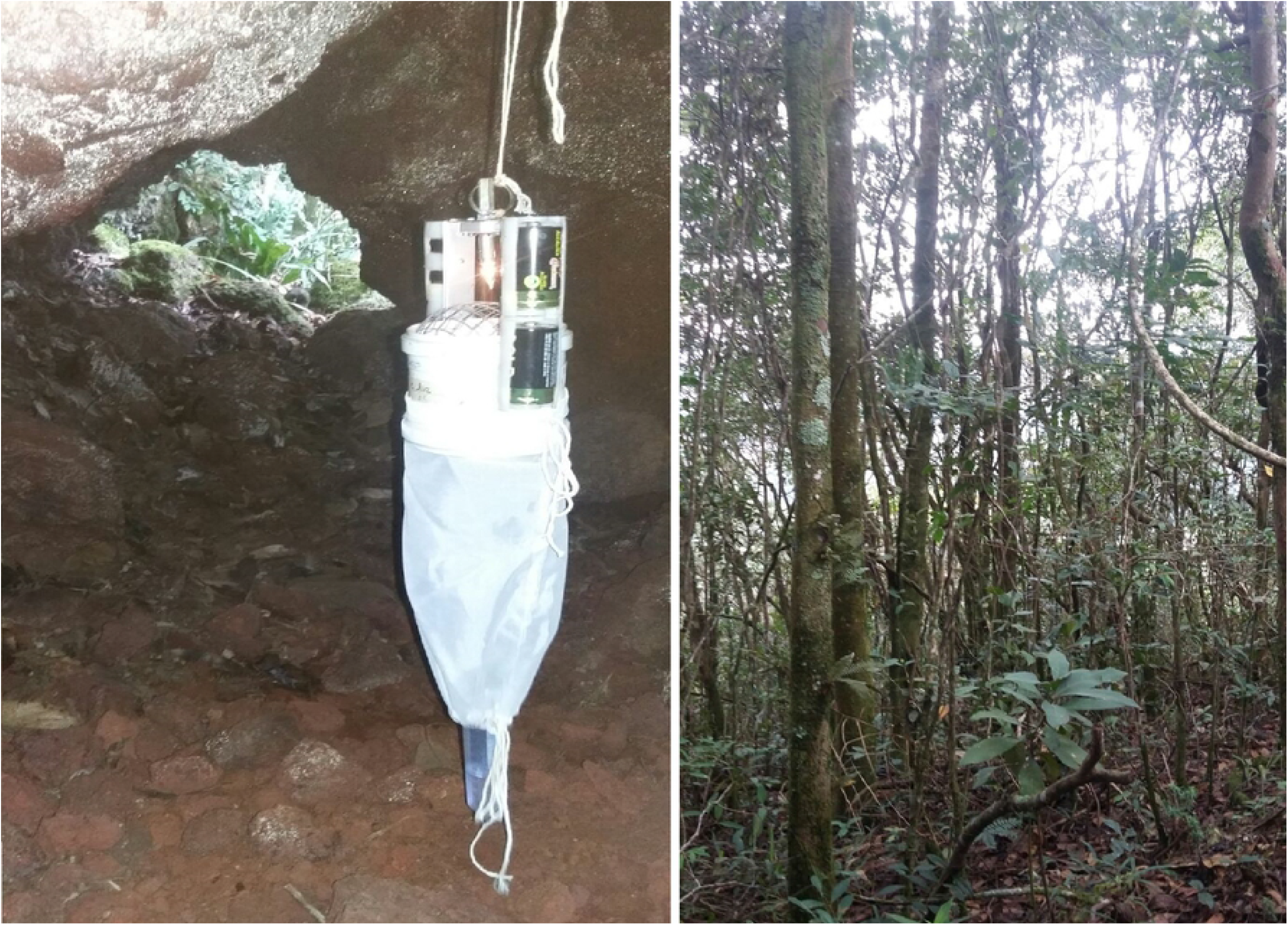
The RM38 cave and their surroundings in Parque Estadual Serra do Rola Moça (PESRM), Minas Gerais, Brazil.

**Fig 4.**
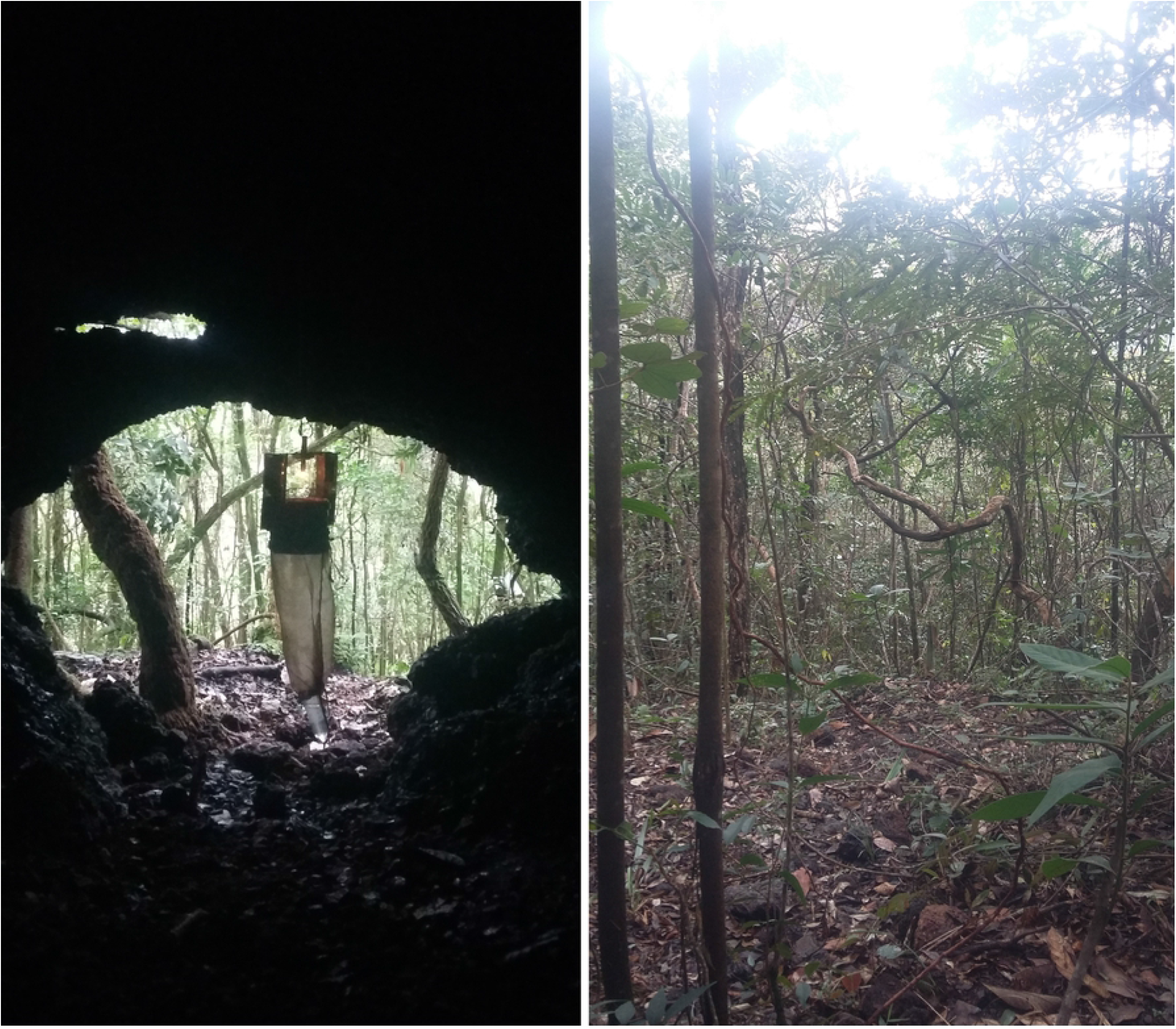
The RM39 cave and their surroundings in Parque Estadual Serra do Rola Moça (PESRM), Minas Gerais, Brazil.

**Fig 5.**
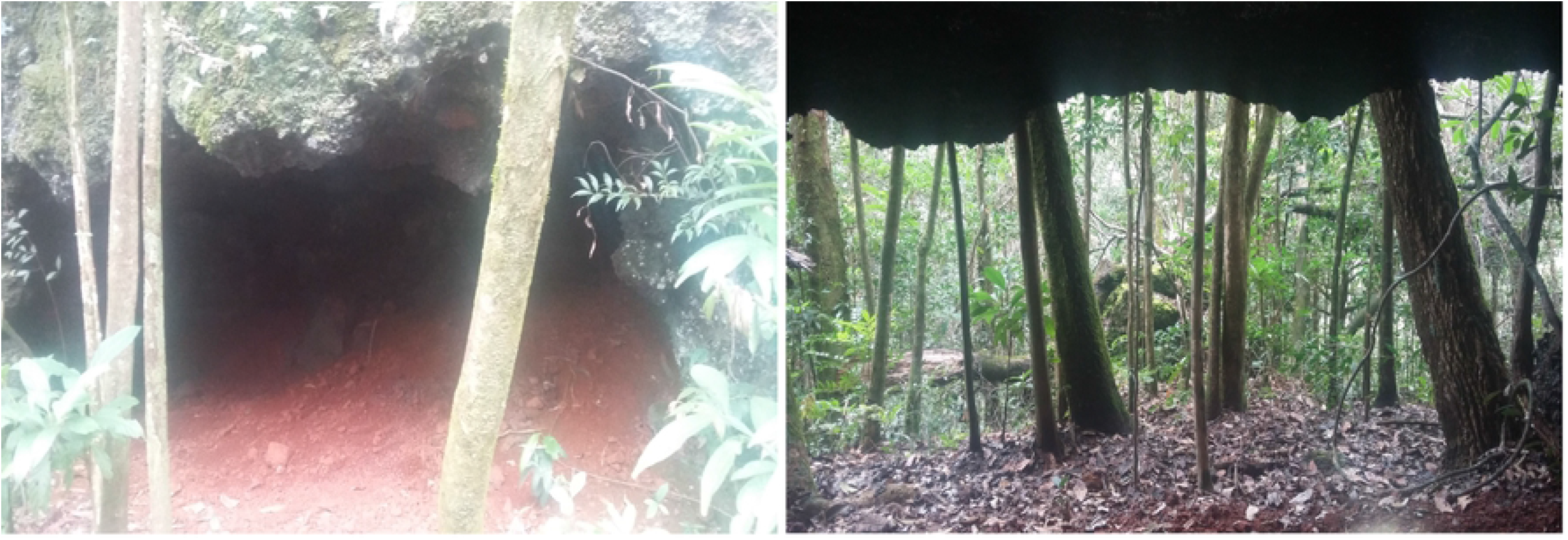
The RM40 cave and their surroundings in Parque Estadual Serra do Rola Moça (PESRM), Minas Gerais, Brazil.

**Table 1.** Description of caves and their surroundings sampled at Moeda Sul (MS) and Parque Estadual Serra do Rola Moça (PESRM), municipality of Nova Lima-MG in the region of the Quadrilátero Ferrífero, from February 2014 to March 2015.

| Locations | Description of caves |  |  |  |
| --- | --- | --- | --- | --- |
|  | Caves names | Coordinates | Elevation (m a.s.l) | Horizontal extension (m) |
| MS | MS Cave | 20°12'7"S, 43°58'2"W | 1,440 | 70 |
|  | RM38 Cave | 20°00'48"S, 43°58'43"W | 1,363 | 50 |
| PESRM | RM39 Cave | 20°00'56"S, 43°58'38"W | 1,272 | 30 |
|  | RM40 Cave | 20°00'59"S, 43°58'40"W | 1,264 | 20 |

The caves located at MS and PESRM are all “canga” caves with no running water and are located in the municipality of Nova Lima in a transition zone between the Cerrado and Atlantic Forest biomes. The cave and their surroundings of MS is located in a region of intense exploitation for real estate and urbanization [24], while the caves and their surroundings of PESRM are located in a largely conserved forest area. Collection of sand flies

The collection of sand flies was performed under the license N° 45636-1 of the Ministério do Meio Ambiente do Brasil and N° 082/2014 of the Instituto Estadual de Florestas de Minas Gerais.

Systematic collections were performed using model HP automatic light traps, [28], throughout an entire year in MS (April 2014 to March 2015) and in PESRM (February 2014 to January 2015). Sampling took place monthly for three consecutive days for a total of 72 h of sampling effort per trap per month.

In place MS, four traps were installed in MS cave and eight in its surroundings for a total of 12 traps. In place PESRM, three traps were placed in RM38 cave, and six traps in its surroundings for a total of nine traps. For the RM39 cave and RM40 cave and their surroundings, two traps were placed in each cave and four to the surroundings of each one of them, total of six traps in each. For the entire study were 33 traps, 11 traps inside of caves and 22 in their surroundings. For all caves and their surroundings sampled, the first trap inside a cave was at its entrance and the other was set every 10 m further into the cave. In the surroundings, the first trap was set in the forest 10 m from the cave entrance and the others were set every 10 m further away from the cave into the forest. The traps were installed approximately 1 m above the ground in all cave and surrounding environments.

The number of traps installed inside each cave was based on the horizontal extension each cave and also the difficulty of moving inside them, making it impossible to use a greater number of traps. Thus, for a better sampling of sand fly fauna at this forest fragments that surrounded these caves we chose to double the number of traps in the surroundings of each cave.

After sampling, the traps were removed and the sand flies killed in glycerinated alcohol, sorted and sexed. Males were then placed in test tubes containing 70% alcohol and females were placed in test tubes containing 6% DMSO for future study of *Leishmania* DNA detection.

### Sand fly identification

Male sand flies were prepared, mounted in Canada Balsam and identified following the classification of [29] and nomenclature of the species of the genus *Psychodopygus* follows the proposal of [30]. For female identification, part of the body (head and the last three segments of the abdomen) was separated and mounted in Berlese medium, with the head positioned ventrally. Specimens will be deposited in the Coleção de Flebotomíneos of the Centro de Pesquisas René Rachou/Fiocruz (FIOCRUZ/COLFLEB).

### Statistical analysis

The similarity of the species composition of sand flies per sampling point of collection among the entrance, interior, and surroundings environments of each sampled cave (MS cave, RM39 cave and RM40 cave) at places MS and PESRM was determined using the Non-Metric Multidimensional Scaling (NMDS) based on a dissimilarity matrix calculated with the Jaccard índex (with regard to the presence and absence of species). The Non-parametric PERMANOVA (Multivariate Analysis of Permutation) with 999 replications across the distance of matrices [31] was used to test if the species composition of sand flies differed among the environments (entrance, interior, and surroundings) of each sampled cave (MS cave, RM39 cave and RM40 cave). All analyzes were performed using the software R 3.2.4 [32] (Fig. 2).

For the NMDS and the Non-parametric PERMANOVA was considered one sampling point at the entrance, interior and surrounding of each cave studied. No phlebotomine sand fly was caught inside RM38 cave, and so this cave and its surroundings were removed from the Non-Metric Multidimensional Scaling (NMDS) analysis.

## Results

A total of 375 phlebotomine sand flies were collected from six genera and 14 species. The total of specimens by sex and the number of species in all environments (cave and surroundings) are shown in Table 2. The most abundant species was *Evandromyia tupynambai* (54.7%), followed by *Brumptomyia troglodytes* (25.6%), *Evandromyia edwardsi* (6.1%), *Psathyromyia brasiliensis* (4.8%) and *Lutzomyia longipalpis* (4.3%). All other species together amounted to 4.5% (Table 2).

**Table 2.**
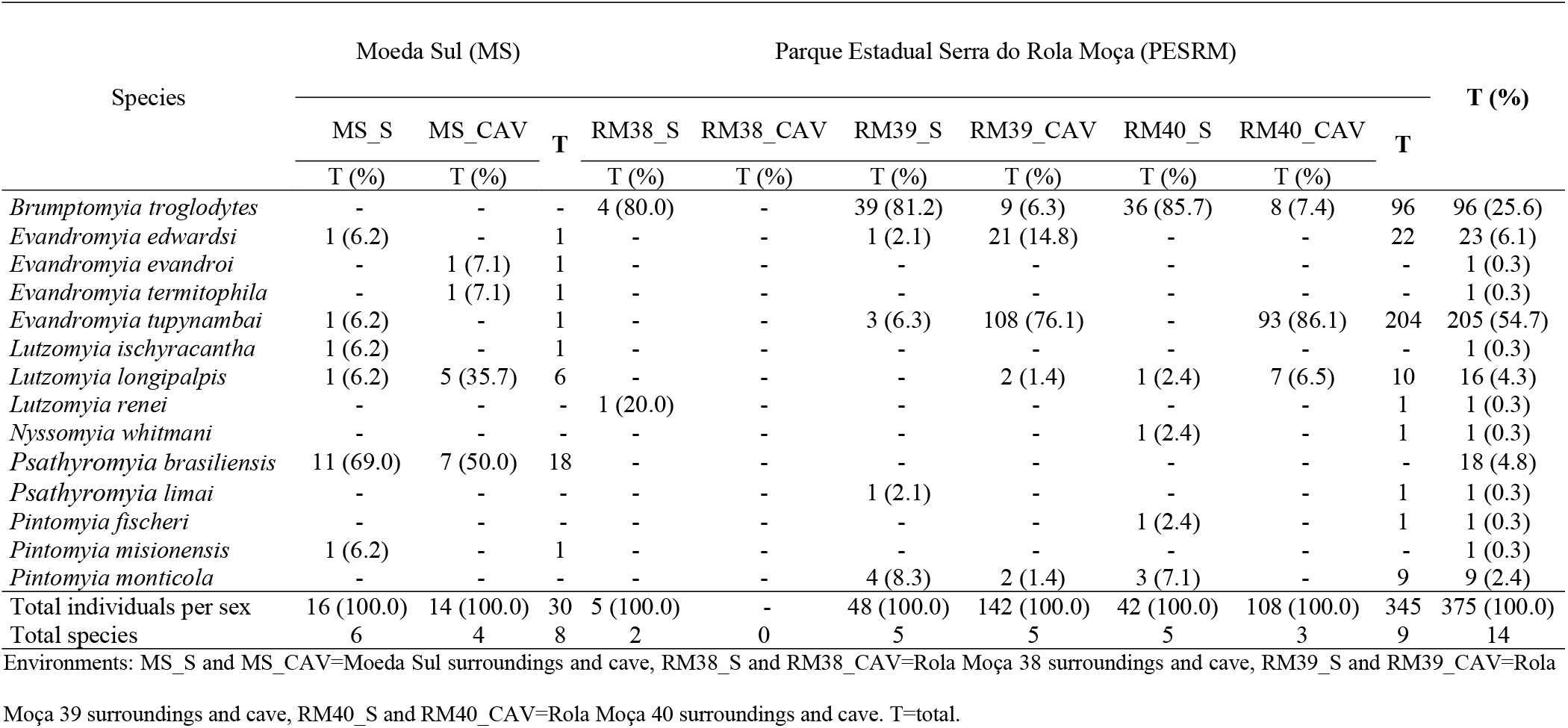
Phlebotomine sand flies collected in caves and their respective surroundings in Moeda Sul (MS) and Parque Estadual Serra do Rola Moça (PESRM), municipalities of Nova Lima-MG, from February 2014 to March 2015. “S” after the cave name indicates samples collected in the area surrounding the cave, and “CAV” indicates samples collected within the cave.

In place MS, 30 individuals were collected, 16 inside the MS cave and 14 from its surroundings, with the most abundant species in the cave being *Pa. brasiliensis* (50%) and *Lu. longipalpis* (35.7%), whereas in the surroundings the most abundant was *Pa. brasiliensis* (69%). In place PESRM, only five individuals were collected from the surroundings of RM38 cave (none from inside the cave); these were *Br. troglodytes* (80.0%) and *Lutzomyia renei* (20.0%). Inside RM39 cave and from its surroundings a total of 190 individuals were collected (48 in the cave and 142 from its surroundings), with *Ev. tupynambai* (76.1%) being the most abundant species in the cave, and *Br. troglodytes* (81.2%) the most abundant in its surroundings. Inside RM40 cave and from its surroundings a total of 150 individuals were collected (42 in the cave and 108 in its surroundings) with *Ev. tupynambai* (86.1%) being the most abundant in the cave and *Br. troglodytes* (85.7%) the most abundant in its surroundings (Table 2).

Non-Metric Multidimensional Scaling (NMDS) calculated with the presence and absence data of the different species of sand flies explained 80% of the variation of the data on two axes (Fig 6). The permutation test showed that the species composition of sand flies did not differ among the entrance, interior, and surroundings environments of each of the caves sampled (MS cave, RM39 cave and RM40 cave) (PERMANOVA: F_1_=0.49; R^2^=0.06; P=0.87).

**Fig 6.**
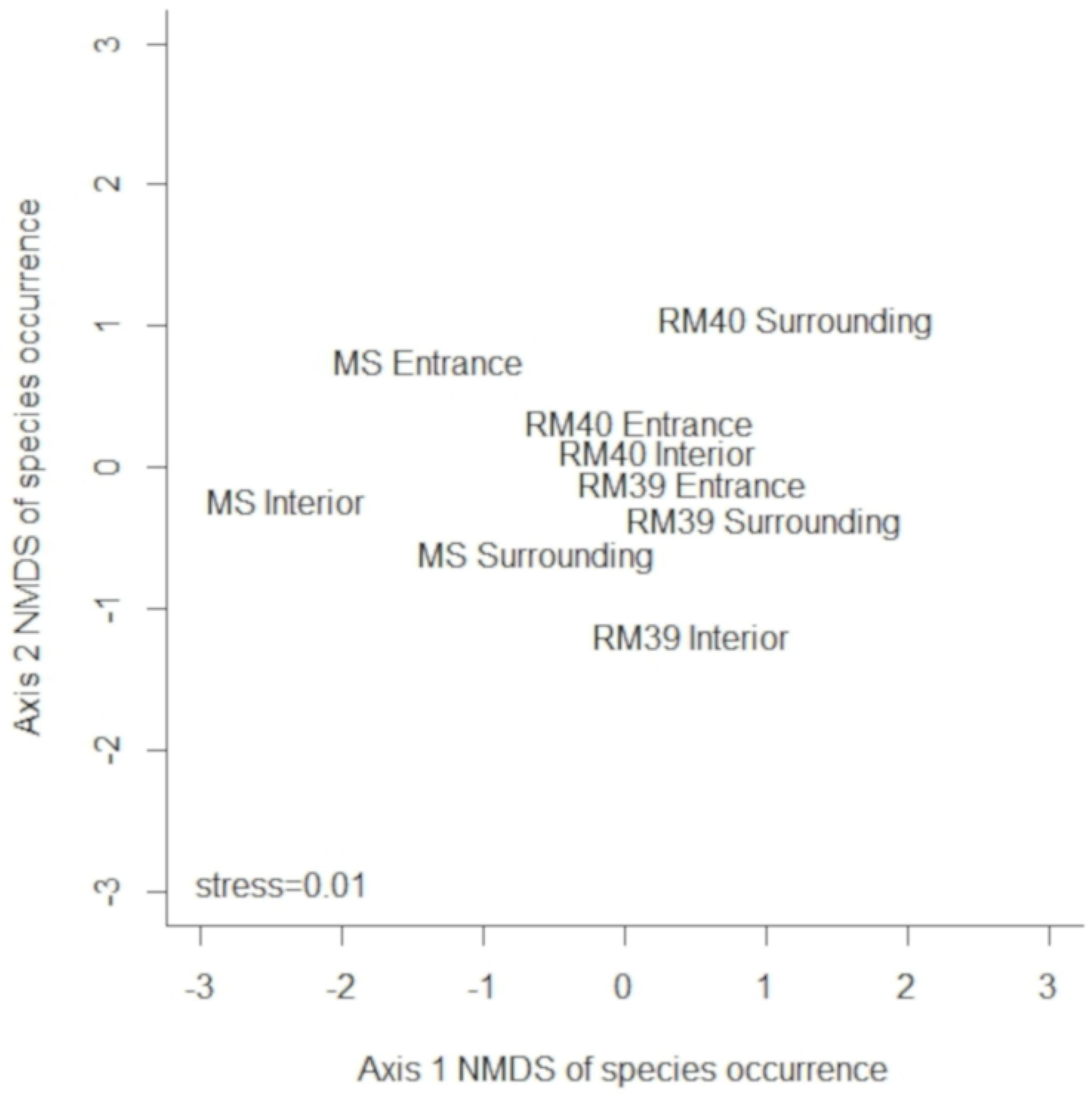
Non-Metric Multidimensional Scaling (NMDS) from a Jaccard dissimilarity matrix calculated with the presence and absence data of the different species of sand flies among the entrance, interior, and surroundings environments of each sampled cave (MS cave, RM39 cave and RM40 cave).

## Discussion

Our results showed that the species composition of sand flies did not differ among the entrance, interior, and surroundings environments of each sampled cave in the two places, MS and PESRM, indicating that it is possible that many species use the caves as places for rest, protection, shelter, breeding, and feeding, as reported by [16]. According to [33], in the ferruginous systems, the external environmental severity is remarkable, and this can lead many species to shelter in the caves, a fact observed with the sand flies community recorded in our study. According to [5], cave environments are commonly used by sand flies for shelter or as breeding sites, and also as their habitat, being the latter also observed by [7], that recorded the presence of *Lutzomyia maruaga,* a species restricted to the cave interior located in the municipality of Manaus-AM.

The phlebotomine sand fly fauna collected inside and outside of the studied caves was very diverse, with species restricted to caves, other species restricted to cave surroundings and others common to both environments. The finding of a rich sand fly fauna in cave environments has been reported previously by [16] in caves in the municipality of Lassance, state of Minas Gerais, [10], in the municipality of Pains, state of Minas Gerais and [19] in caves in the state of São Paulo.

The most abundant species in our study, *Br. troglodytes, Ev. edwardsi, Ev. tupynambai* and *Lu. longipalpis,* are commonly found inside and outside the caves [4, 34, 35, 8, 16, 10]. Other species abundant was *Pa. braziliensis,* species with few records in caves.

The fact that the caves and its surroundings of PESRM are in a conserved forest fragment, explains the occurrence of the species *Ev. tupynambai* and *Br. trogodytes* as the most abundant inside and outside of the caves, respectively, at that location. These species are very common in conserved forest environments, although they have already been found in shelters of domestic animals, marginal areas and human habitations [36, 34, 15, 37]. *Evandromyia tupynambai* as the most abundant species inside the caves was a surprise, because despite its wide distribution, no information in the literature about the presence of this species in cave environment was recorded, but a very close species *(Evandromyia petropolitana)* was described in a stone slot [38]. However, the record of *Br. trogodytes* as the most abundant species in the cave surroundings was to be expected since it is a characteristic species of forest areas [36].

The cave and its surrounding of MS is situated in a very anthropized region surrounded by a residential condominium and a BR, therefore, it was not surprising the occurrence of the species *Pa. brasiliensis* e *Lu. longipalpis* as the most abundant since they are two very common species in areas of secondary forest, urban areas and shelters of domestic animals [39, 9, 10]. The finding of *Pa. brasiliensis* as the first most abundant species inside the cave has no record in the literature, but to have been more abundant in the cave surrounding does not seem unusual, since it is a species that occurs in anthropized areas [39]. *Lutzomyia Longipalpis* was also one of the most abundant species inside the cave, corroborating [10] that recorded this species as one of the most abundant inside the caves in other area of Minas Gerais State.

The species incriminated in the transmission of leishmaniasis found in this study, *Lu*. *longipalpis* and *Nyssomyia whitmani*, have also been recorded in cave environments by other authors [4, 5, 35, 16, 10].

The biggest part of the sand flies collected are trogloxenes and the species *Ev. tupynambai* and *Ev. edwardsi,* the two most abundant species, can be considered troglophiles, since they were recorded in almost every month of collection with more than 90% of the individuals registered inside the cave. This result corroborates with [16] who reported the presence of trogloxenes and troglophiles phlebotomines.

An interesting fact of the present study was that no sand fly was recorded at the entrance and the interior of RM38 cave located in the PESRM, only in its surroundings. One possible explanation would be the very small entrance of this cave in relation to the other caves of PESRM. [4] studying several caves in the Serra da Bodoquena (Mato Grosso do Sul) observed for three caves with low densities of recorded sand flies, that the cave with the highest density of the three was with a wide entrance which may be more attractive for several trogloxenes species.

In summary, a rich phlebotomine fauna was recorded with sand fly communities similar among the entrance, interior, and surroundings environments of each sampled cave at places, MS and PESRM. Furthermore, species incriminated in *Leishmania* transmission were recorded, and the presence of *Lu. Longipalpis* in the surrounding of cave MS, even if in low abundance, deserves special attention, for being a close location to a residential condominium, since this species is the main vector of *Leishmania infatum,* causative agent of visceral leishmaniasis [40]. Therefore, our suggest results that both caves and their surroundings are important for the maintenance of their sand fly communities.

## Acknowledgments

We thank Fundação de Amparo à Pesquisa do Estado de Minas Gerais/FAPEMIG (RDP-00149-10, PPM-00438-14) FAPEMIG, Conselho Nacional de Pesquisas/CNPq (N° 301421/2013-7) CNPq, and VALE S.A for funding and a research grant, and the Coordination of Training of Upper-Level Personnel (CAPES) for the awarded grant. We also thank D. Capucci for editing the map of study area that is shown in Fig 1.

